# Environmental stability of SARS-CoV-2 on different types of surfaces under indoor and seasonal climate conditions

**DOI:** 10.1101/2020.08.30.274241

**Authors:** Taeyong Kwon, Natasha N. Gaudreault, Juergen A. Richt

## Abstract

We report the stability of SARS-CoV-2 on various surfaces under indoor, summer and spring/fall conditions. The virus was more stable under the spring/fall condition with virus half-lives ranging from 17.11 to 31.82 hours, whereas under indoor and summer conditions the virus half-lives were 3.5–11.33 and 2.54–5.58 hours, respectively.

## The Study

Severe acute respiratory coronavirus 2 (SARS-CoV-2) which first emerged in a wet market in Wuhan, China, is responsible for the current pandemic. Although transmission of SARS-CoV-2 mainly occurs through infectious droplets or close contact with an infected person, the virus droplet can survive and remain infectious on inanimate surfaces, which can contribute to the spread of the virus (1). Previous studies showed that virus remained infectious from hours to days on various type of surfaces under various temperature-controlled environmental conditions (2–4). However, virus stability on surfaces under different climate conditions which could be used to predict seasonality of SARS-CoV-2, is poorly understood. In this manuscript, we evaluated the stability of SARS-CoV-2 on different types of surfaces under indoor, summer and spring/fall conditions to estimate the biological half-life of the virus.

We tested SARS-CoV-2 stability on 12 material surfaces including nitrile glove, Tyvek, N95 mask, cloth, Styrofoam, cardboard, concrete, rubber, glass, polypropylene, stainless steel and galvanized steel (see Technical Appendix). Each material surface was placed in a 6-well or 12-well plate and 50 μl of virus inoculum consisting of 5×10^4^ TCID_50_ SARS-CoV-2 (strain USA-WA1/2020) in DMEM with 5% FBS was added onto each material. The positive control had the same amount of virus in medium in a sealed 2mL tube. The virus was air-dried inside a biosafety cabinet (approximately 4.5 hours). The plate with the virus-contaminated material was incubated under three different conditions: 21°C/60% relative humidity (RH), 25°C/70% RH and 13°C/66% RH, environmental conditions simulating indoor setting, summer, and spring/fall conditions for the Midwestern U.S., respectively (Technical Appendix Table 1). At each time point indicated, infectious virus was recovered in 2 mL media through vigorous vortexing for 10 seconds. Cardboard was soaked with media for 5 minutes and vortexed for 10 seconds. The recovered virus was titrated on Vero E6 cells and virus titer was calculated by the Reed-Muench method. The assay was performed in triplicate. A best-fitting line was estimated using a linear regression model in order to calculate the virus half-life on each surface as a −log_10_(2)/slope and tested for statistical significance using default analysis which is compatible to analysis of covariance in GraphPad Prism 5.

SARS-CoV-2 was relatively stable in medium throughout the study phase, showing a 1.17-log reduction of virus titer at 96 hours post-contamination (hpc) at 25°C/70% RH (Figure 1). We found a 1-log reduction of virus after 4.5 hours at room temperature (21°C/60% RH) on all materials (10^3.3^ to 10^4.2^ TCID_50_), except for cloth (10^2.4^ to 10^2.7^ TCID_50_), which served as the starting titers for the linear regression model. At 21°C/60% RH, infectious virus was recovered from cloth up to 24 hpc, from concrete, polypropylene, stainless steel and galvanized steel up to 72 hpc, and from nitrile gloves, Tyvek, N95 mask, Styrofoam, cardboard, rubber and glass up to 96 hpc. In contrast, viable virus disappeared quickly under summer conditions (25°C/70% RH) and was undetectable on cloth, cardboard, concrete and stainless steel at 48 hpc, and on nitrile gloves, Tyvek, N95 mask, Styrofoam, rubber, glass, polypropylene, galvanized steel at 72 hpc. However, we observed longer survival times at spring/fall conditions (13°C/66% RH). Virus titers on surfaces ranged from 10^1.1^ to 10^2.3^ TCID_50_ at 168 hpc, except for cloth with virus only detectable up to 72 hpc. Half-lives of SARS-CoV-2 on surfaces ranged from 3.5 to 12.86 hours at 21°C/60% RH, 2.54 to 5.58 hours at 25°C/70% RH, and 17.11 to 31.82 hours at 13°C/66% RH (Table 1). The virus survived significantly longer on all surfaces at spring/fall conditions (13°C/66% RH) when compared to summer and indoor conditions. Similarly, we found a significant difference in virus survival on surfaces between indoor and summer conditions except for cloth.

**Figure 1.**
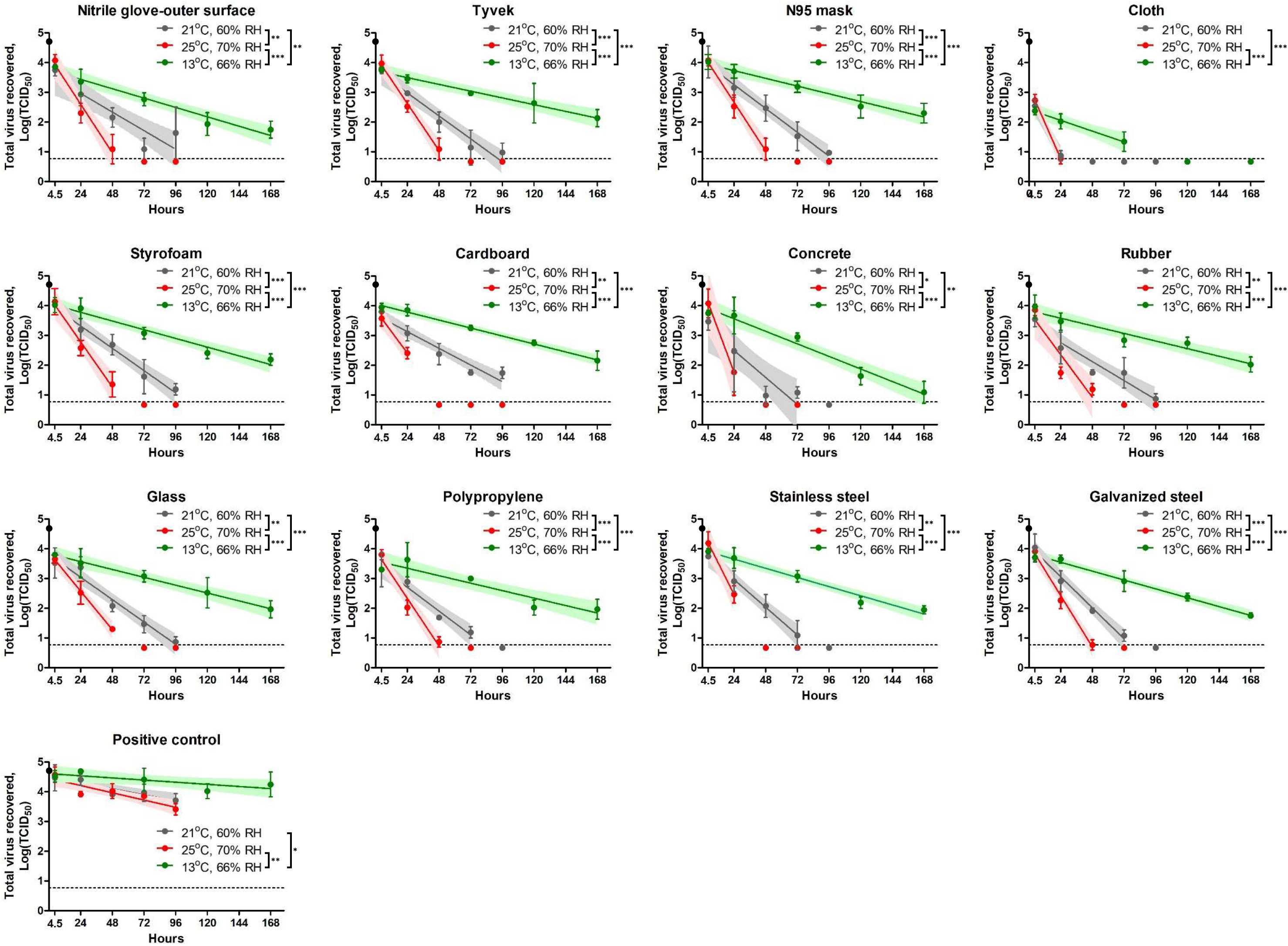
Stability of severe acute respiratory coronavirus 2 (SARS-CoV-2) on different types of surfaces. Each figure represents the virus decay on each surface. Total 50 μl of virus inoculum (5×10^4^ TCID_50_, black dot) was added onto each material and dried for 4.5 hours inside a biosafety cabinet. The virus survival was evaluated under three different conditions: at 21°C/60% RH (grey), 25°C/70% RH (red) and 13°C/66% RH (green). The infectious virus was recovered at 4.5 (after drying period), 24, 48, 72, and 96 hours post-contamination (hpc) at 21°C/60% RH and 25°C/70% RH and 4.5, 24, 72, 120, and 168 hpc at 13°C/66% RH. Virus titer at each time point was expressed as mean log_10_ transformed titer with standard deviation. Linear regression models were estimated; the solid line and its shade area represent an estimated best fit model and 95% confidence intervals, respectively. Limit of detection (LOD) in each titration assay was 10^0.968^ TCID_50_ and a negative result is represented as a half value of LOD, 10^0.667^ TCID_50_. The dash line shows LOD in triplicate, 10^0.767^ TCID_50_, when there was LOD in one replicate, but negative in two other replicates. Statistical significance between two slopes of linear regression models is represented as * (*p* < 0.05), ** (*p* < 0.01), *** (*p* < 0.001).

**Table 1.** Half-lives of severe acute respiratory coronavirus 2 (SARS-CoV-2) on different types of surfaces. The virus decay rates were evaluated under three different conditions, 21°C/60% RH, 25°C/70% RH and 13°C/66% RH, which simulate indoor, summer and spring/fall conditions, respectively.

| Surface materials | 21°C, 60% relative humidity<br>(indoor condition) |  |  | 25°C, 70% relative humidity<br>(Summer condition) |  |  | 13°C, 66% relative humidity<br>(Spring/fall condition) |  |  |
| --- | --- | --- | --- | --- | --- | --- | --- | --- | --- |
| | Half-life<br>(hours) | 95%<br>confidence<br>interval<br>(hours) | $r^2$ | Half-life<br>(hours) | 95%<br>confidence<br>interval<br>(hours) | $r^2$ | Half-life<br>(hours) | 95%<br>confidence<br>interval<br>(hours) | $r^2$ |
| Nitrile gloves –<br>outer surface | 11.56 | 8.27, 19.21 | 0.69 | 4.42 | 3.5, 6.03 | 0.92 | 22.94 | 18.73, 29.63 | 0.88 |
| Tyvek | 9.36 | 7.76, 11.79 | 0.89 | 4.57 | 3.84, 5.63 | 0.96 | 31.82 | 24.65, 44.82 | 0.81 |
| N95 mask | 9.01 | 7.57, 11.12 | 0.91 | 4.4 | 3.64, 5.57 | 0.95 | 27.77 | 22.5, 36.27 | 0.87 |
| Cloth | 3.5 | 2.77, 4.75 | 0.97 | 2.99 | 2.45, 3.84 | 0.98 | 19.94 | 13.94, 34.95 | 0.81 |
| Styrofoam | 9.62 | 8.04, 11.98 | 0.9 | 4.75 | 3.73, 6.53 | 0.92 | 24.67 | 20.6, 30.73 | 0.9 |
| Cardboard | 12.86 | 10.52, 16.54 | 0.88 | 5.03 | 3.5, 8.95 | 0.91 | 26.93 | 23.55, 31.42 | 0.95 |
| Concrete | 7.96 | 5.25, 16.44 | 0.65 | 2.54 | 1.55, 6.98 | 0.83 | 17.11 | 14.38, 21.14 | 0.91 |
| Rubber | 11.33 | 8.95, 15.45 | 0.83 | 5.03 | 3.63, 8.18 | 0.84 | 28.27 | 22.4, 38.32 | 0.84 |
| Glass | 9.6 | 8.05, 11.89 | 0.91 | 5.58 | 4.72, 6.82 | 0.96 | 27.34 | 21.72, 36.87 | 0.84 |
| Polypropylene | 9.02 | 7.22, 12.03 | 0.89 | 4.51 | 3.74, 5.68 | 0.95 | 28.75 | 21.52, 43.36 | 0.76 |
| Stainless | 7.75 | 6.39, 9.86 | 0.92 | 3.41 | 2.36, 6.16 | 0.91 | 23.46 | 20.16, 28.08 | 0.93 |
| Galvanized | 6.93 | 5.88, 8.43 | 0.94 | 4.19 | 3.68, 4.85 | 0.98 | 24.22 | 21.3, 28.08 | 0.95 |
| Positive control | 35.54 | 23.19, 75.88 | 0.56 | 29.48 | 20.85, 50.39 | 0.68 | 100.68 | 52.35, 1346.89 | 0.3 |

Potential modes of transmission of SARS-CoV-2 include direct contact with an infected person via droplets, inhalation of aerosol or infectious body fluids, and exposure to contaminated surfaces (fomite). To date, there is no scientific report which demonstrates SARS-CoV-2 infection via contaminated surfaces. However, the role of fomites in transmission of SARS-CoV-2 is debated because the virus has been detected on environmental surfaces as well as personal protective equipment in hospitals and households (5, 6). In addition, indirect transmission of SARS-CoV-2 has been supported by a cluster of SARS-CoV-2 infection cases in a shopping mall, in which contact tracing failed to find any evidence for direct contact to an infected person, only to sharing of facilities (7). In this respect, our study highlights the possible role of contaminated surfaces in SARS-CoV-2 transmissions because SARS-CoV-2 remained viable and infectious on surfaces for 1 to 4 days at indoor conditions (21°C/60% RH), 1 to 3 days during summer conditions (25°C/70% RH) and over 7 days during spring/fall conditions (13°C/66% RH).

Van Doremalen et al. (3) described that the SARS-CoV-2 half-life which ranges from 3.46 to 6.81 hours on cardboard, plastic and stainless steel at 22°C/40% RH. Chin et al (2) reported a half-life of 4.8 to 23.9 hours on glass, banknotes, inner and outer mask layers, polypropylene and stainless steel at 22°C/65% RH. We found the half-life on most surfaces at 21°C/60% RH is 6.93–12.86, but the virus is quickly inactivated on cloth with a 3.5 hours half-life. The difference might be explained by the composition of the virus inoculum (e.g., FBS concentration), the volume of inoculum, different preparation of material and the different environmental conditions. However, our results, along with other two studies, showed that SARS-CoV-2 is able to survive on some surfaces for several days under indoor conditions, which might play a potential role in virus transmission. The longest half-life of the virus was found in spring/fall conditions (13°C/66% RH), followed by indoor conditions (21°C/60% RH) and summer conditions (25°C/70% RH); this suggests that virus stability on surfaces is highly dependent on temperature and RH. Prolonged virus survival on surfaces in spring/fall and winter might support SARS-CoV-2 transmission through contaminated fomites and potentially contribute to new outbreaks and/or seasonal occurrence in the post-pandemic era, a scenario described for influenza virus and other human coronaviruses (8).

Our study showed a remarkable persistence of infectious SARS-CoV-2 on various types of surfaces, especially under spring/fall climate conditions. However, virus stability was highly dependent on the substrate as well as temperature and humidity. Previous studies showed reduced virus stability in human nasal mucus and sputum when compared to culture medium (9) even at 4°C/40% RH, whereas addition of bovine serum albumin into the virus inoculum increased SARS-CoV-2 survival times (10). In addition, exposure to simulated sunlight accelerated the inactivation of the virus on stainless steel (11), indicating that additional factors play a role in SARS-CoV-2 survival on surfaces in field settings.

In conclusion, our study determines the half-life of SARS-CoV-2 on diverse surfaces under different climatic conditions, which correlates to the potential risk of contaminated surfaces to spread the virus. It clearly demonstrates, that the virus survives longer under spring/fall not summer conditions. Therefore, practice of good personal hygiene and regular disinfection of potentially contaminated surfaces remains a critical tool to minimize the risk of infection through contaminated surfaces.

## Acknowledgements

Funding for this study was provided through grants from the National Bio and Agro-Defense Facility (NBAF) Transition Fund, Kansas State University internal funds, and the NIAID Centers of Excellence for Influenza Research and Surveillance under contract number HHSN 272201400006C to JAR.

## Biographical Sketch

Mr. Kwon is a Doctor of Veterinary Medicine and presently a PhD student at Kansas State University. His research interest are transboundary animal disease and emerging zoonotic diseases.

## Technical appendix

## Preparation of surface materials

Materials used in this study were nitrile glove (Kimberly-Clark Professional™ Kimtech™ G3 Sterile Sterling™ Nitrile Gloves), Tyvek (DuPont™ Tyvek IsoClean Sleeves. Clean Processed & Sterile, White), N95 mask (3M N95 mask 1870), cloth (65% polyester and 35% cotton from local source), styrofoam (50mL centrifuge tube-foam rack, CELLTREAT Scientific Products), cardboard (inner packing, TPP T75 flask), concrete (Fast-setting concrete mix, The Home Depot), rubber (The Home Depot), glass (Electron Microscopy Sciences), polypropylene (biohazard autoclave bag, ThermoFisher), stainless steel (Metal Remnant Inc.), and galvanized steel (The Home Depot). Materials were cut into small pieces, washed, dried and autoclaved (depending on material). To make concrete, the coarse aggregate was removed by a strainer, and the fine aggregate was mixed with water according to the manufacturer’s instruction. Mixture was poured into a silicone mold and air-dried in biosafety cabinet overnight.

## U.S. Midwest climate conditions

Maximum and minimum temperature and relative humidity (RH) data at Manhattan, Kansas, was acquired from National Service Forecast Office on 5/11/2020 (https://w2.weather.gov/climate/index.php?wfo=top). Average temperature and RH was calculated for each season. Climate conditions for spring and fall were combined since their average temperature and RH were similar. Spring/fall and summer conditions were 13°C/66% RH and 25°C/70% RH, respectively.

---

**Technical Appendix Table 1.**
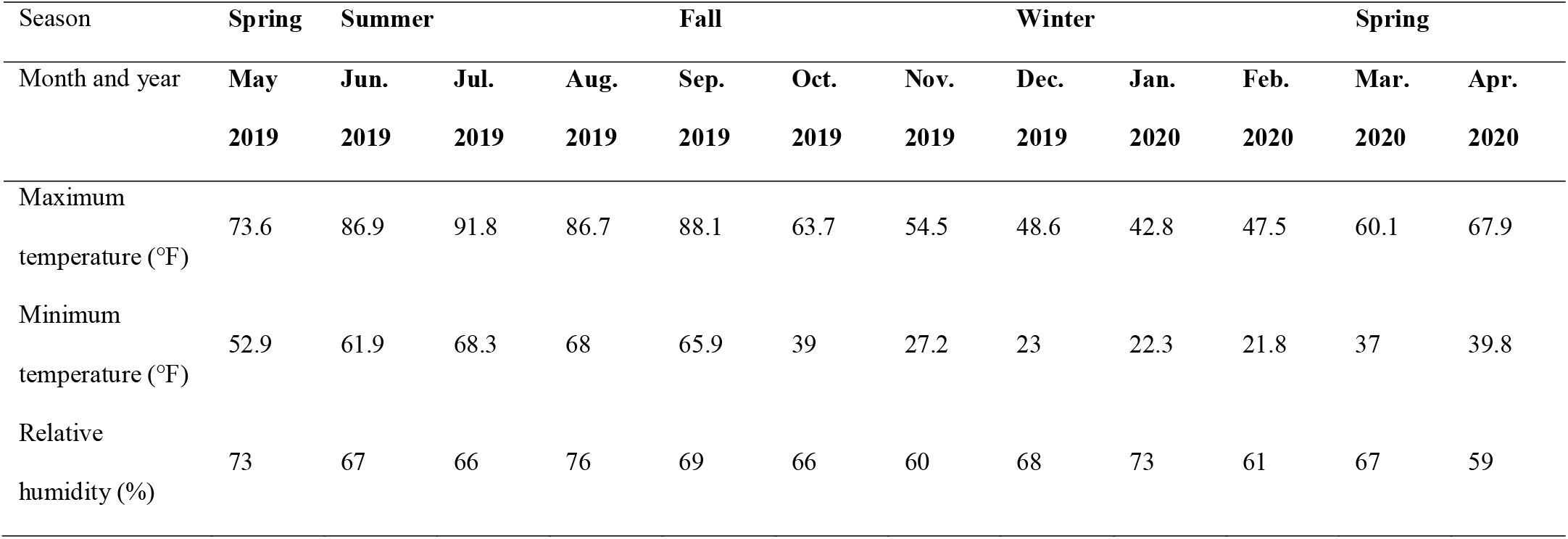
Maximum and minimum temperature and relative humidity data for Manhattan, Kansas

## References

1. World Health Organization. Transmission of SARS-CoV-2: implications for infection prevention precautions. 2020 [Cited 2020 Aug 12] https://www.who.int/news-room/commentaries/detail/transmission-of-sars-cov-2-implications-for-infection-prevention-precautions.

2. Chin AWH, Chu JTS, Perera MRA, Hui KPY, Yen H-L, Chan MCW, et al. Stability of SARS-CoV-2 in different environmental conditions. The Lancet Microbe. 2020;1(1):e10.

3. van Doremalen N, Bushmaker T, Morris DH, Holbrook MG, Gamble A, Williamson BN, et al. Aerosol and surface stability of SARS-CoV-2 as compared with SARS-CoV-1. N Engl J Med. 2020;382(16):1564–7.

4. Kratzel A, Steiner S, Todt D, V’Kovski P, Brueggemann Y, Steinmann J, et al. Temperature-dependent surface stability of SARS-CoV-2. J Infect. 2020;81(3):474–6.

5. Ong SWX, Tan YK, Chia PY, Lee TH, Ng OT, Wong MSY, et al. Air, surface environmental, and personal protective equipment contamination by severe acute respiratory syndrome coronavirus 2 (SARS-CoV-2) from a symptomatic patient. JAMA. 2020;323(16):1610–12.

6. Jiang XL, Zhang XL, Zhao XN, Li CB, Lei J, Kou ZQ, et al. Transmission potential of asymptomatic and paucisymptomatic severe acute respiratory syndrome coronavirus 2 infections: A 3-family cluster study in China. J Infect Dis. 2020;221(12):1948–52.

7. Cai J, Sun W, Huang J, Gamber M, Wu J, He G. Indirect virus transmission in cluster of COVID-19 cases, Wenzhou, China, 2020. Emerg Infect Dis. 2020;26(6):1343–5.

8. Kissler SM, Tedijanto C, Goldstein E, Grad YH, Lipsitch M. Projecting the transmission dynamics of SARS-CoV-2 through the postpandemic period. Science. 2020;368(6493):860–8.

9. Matson MJ, Yinda CK, Seifert SN, Bushmaker T, Fischer RJ, van Doremalen N, et al. Effect of environmental conditions on SARS-CoV-2 stability in human nasal mucus and sputum. Emerg Infect Dis. 2020;26(9).

10. Pastorino B, Touret F, Gilles M, de Lamballerie X, Charrel RN. Prolonged infectivity of SARS-CoV-2 in fomites. Emerg Infect Dis. 2020;26(9).

11. Ratnesar-Shumate S, Williams G, Green B, Krause M, Holland B, Wood S, et al. Simulated sunlight rapidly inactivates SARS-CoV-2 on surfaces. J Infect Dis. 2020;222(2):214–22.

